# Deletion of absent in melanoma (AIM) 2 gene alters bone morphology

**DOI:** 10.1101/2024.01.05.574199

**Authors:** Zhenwei Gong, Manisha Dixit, Sher Bahadur Poudel, Gozde Yildirim, Shoshanna Yakar, Radhika Muzumdar

**Affiliations:** Division of Endocrinology and Diabetes, University of Pittsburgh School of Medicine, Children’s Hospital of Pittsburgh, Pittsburgh, PA 15224; David B. Kriser Dental Center, Department of Molecular Pathobiology, New York University College of Dentistry, New York, NY

**Keywords:** AIM2, Inflammasome, Bone, Bone marrow, Interferon

## Abstract

Absent in Melanoma (AIM) 2 is a gene that is induced by interferon and acts as a cytosolic sensor for double-stranded (ds) DNA. It forms the AIM2 inflammasome, leading to the production of interleukin (IL)-1β and IL-18. Our previous research demonstrated that mice lacking AIM2 exhibit spontaneous obesity, insulin resistance, and inflammation in adipose tissue. In this study, we aimed to explore the impact of AIM2 gene deletion on bone structure in adult and aged mice. Utilizing micro-computed tomography (micro-CT), we discovered that female mice lacking AIM2 showed an increase in the total cross-sectional area at 5 months of age, accompanied by an increase in cortical thickness in the mid-diaphysis of the femur at both 5 and 15 months of age. At 15 months of age, the cortical bone mineral density (BMD) significantly decreased in AIM2 null females compared to wild-type (WT) mice. In AIM2 null mice, both trabecular bone volume and BMD at the distal metaphysis of the femur significantly decreased at 5 and 15 months of age. Similarly, micro-CT analysis of the L4 vertebra revealed significant decreases in trabecular bone volume and BMD in aged AIM2 null females compared to WT mice. Histological examination of femurs from aged mice demonstrated increased bone marrow adiposity in AIM2 null mice, accompanied by a significant increase in CD45-/CD31-/Sca1+/Pdgfa+ adipose progenitor cells, and a decrease in the ratio of CD31-/CD31+ osteogenic progenitor cells, as determined by flow cytometry of bone marrow cells. Our findings suggest that AIM2 deficiency affects bone health by promoting adipogenesis in bone marrow cells and inducing a pro-inflammatory environment, potentially contributing to the decreased bone mineral density.

## INTRODUCTION

Osteoporosis is a prevalent age-related bone disease worldwide, characterized by low bone mineral density, reduced bone mass, and weakened strength, leading to an increased risk of fractures. According to the CDC, the age-adjusted prevalence of osteoporosis among adults aged 50 and older in the United States was 12.6% in 2017-2018, with higher rates in women compared to men (19.6% vs. 4.4%). Management of osteoporosis involves strategies such as calcium and vitamin D intake, exercise, lifestyle changes, and pharmacological therapies including antiresorptive agents, hormonal therapies, and parathyroid hormone analogues [1].

Absent in Melanoma (AIM) 2 is an innate immunity protein belonging to the HIN-200 domain family. It acts as a cytosolic sensor for double-stranded DNA (dsDNA) and forms the AIM2 inflammasome upon binding to dsDNA [2, 3]. Notably, AIM2 functions as a tumor suppressor, and defects or deletion of the AIM2 gene have been linked to early onset and progression of various cancers in both human and murine models [4-6]. While the AIM2 inflammasome has been implicated in inflammatory and infectious diseases like systemic lupus [7], chronic obstructive pulmonary disease [8], and neurodegenerative diseases [9], an inflammasome-independent role of AIM2 has also been identified. Research has shown that AIM2 suppresses colon tumorigenesis through the activation of the DNA-dependent protein kinase (DNA-PK) and protein kinase B/Akt signaling pathway [5].

In our recent study, we reported an inflammasome-independent role of AIM2 in energy homeostasis [10]. AIM2 null mice exhibited an accelerated aging phenotype, characterized by early onset (3 months of age) obesity and insulin resistance [10]. Notably, we observed increased monocyte infiltration in the white adipose tissue (WAT) prior to the onset of obesity, along with elevated M1 macrophages and inflammation at later stages. At the tissue level, we observed increased inflammation, as evident by the increased expression levels of pro-inflammatory cytokines such as TNFα and IL-6 [10]. RNA sequencing analysis of white adipose tissue (WAT) from AIM2 null mice revealed a significant upregulation of inflammation-related genes and interferon-stimulated genes (ISGs). Among these, Ifi202b, an ISG, showed consistent upregulation in multiple tissues of AIM2 null mice [10]. These findings align with previous research demonstrating increased ifi202b expression in splenocytes from AIM2 null mice through upregulated type 1 interferon signaling [11]. Studies have shown that increased ifi202b expression stimulates adipogenesis, and overexpression of ifi202b in mice induces obesity through increased hypertrophy and decreased thermogenesis [12]. Additionally, increased expression of IFI16, the human ortholog of ifi202b, has been observed in the visceral adipose tissue of obese individuals, and ifi16 has been associated with senescence in human fibroblasts [12, 13]. This is consistent with our study showing an association of increased IFI16 expression and overweight and obesity in children [10].

Given the fact that both increased inflammation and adiposity play key roles in bone health, the aim of this study was to investigate the impact of AIM2 deletion on the bone marrow microenvironment and its subsequent effects on bone morphology and mineral density.

## RESULTS

### Absence of AIM2 is associated with impaired cortical bone adaptation

Female AIM2 null mice exhibit a consistent increase in body weight throughout their lifespan (**Fig.1A**), which is primarily attributed to elevated body adiposity [10]. To understand the impact of these changes in body composition on skeletal integrity, we conducted a comprehensive analysis of femurs obtained from 5 and 15-month-old female wild-type (WT) and AIM2 null mice. We observed an enlargement in the radial dimensions of AIM2 null femurs compared to WT femurs. This was evident through a 20% increase in total cross-sectional area (T.Ar) (**Fig.1B**) and a 24% increase in bone area (B.Ar) (**Fig.1C**), and a 10% increase in the marrow area (M.Ar) (**Fig.1D**) at 5 months of age. Additionally, cortical bone thickness showed a 15% increase in AIM2 null females at 5 months of age (**Fig.1E**). The increased T.Ar in AIM2 null mice prompted us to assess bone robustness, calculated as the ratio of radial bone size to bone length (T.Ar/Length) (**Fig.1F**). At 5 months, AIM2 null mice displayed greater bone robustness compared to WT. However, this effect was lost at 15 months, suggesting a possible inhibition of radial expansion in AIM2 null mice. We examined the skeletal adaptation of AIM2 null mice to their increased body weight by plotting bone robustness relative to body weight. In WT mice, a linear and positive relationship between bone robustness and body weight was observed, whereas AIM2 null mice did not exhibit this relationship (**Fig.1G**). Lastly, despite a 3% increase in cortical tissue mineral density (TMD) at 5 months of age, a marked reduction of 8% in TMD was observed during aging in AIM2 null mice (**Fig.1H**).

**Figure 1:**
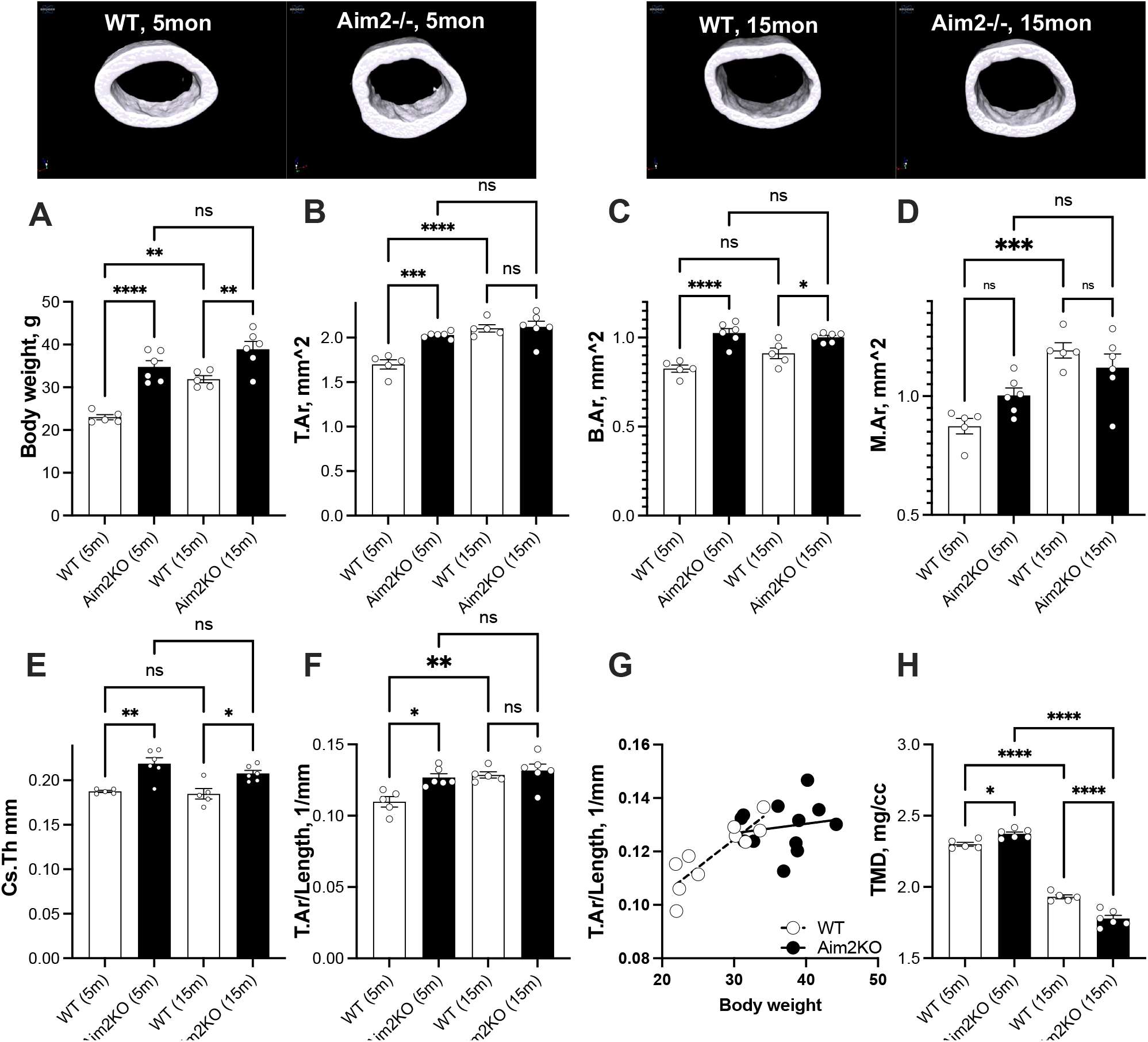
AIM2 null female mice exhibit altered cortical bone morphology. (A) Body weight of WT and Aim2KO mice at 5 and 15 months of age. Cortical bone morphology was assessed at the femur mid-diaphysis, including (B) total cross-sectional area (T.Ar), (C) bone area (B.Ar), (D) marrow area (M.Ar), and (E) cortical bone thickness (Cs.Th). (F) Bone robustness was calculated as the ratio between T.Ar and bone length (T.Ar/Length). (G) Bone robustness was corrected to body weight of 5- and 15-months old mice, showing linear relationship in WT but not in Aim2KO mice. (H) Cortical bone tissue mineral density ((TMD). Significance was determined by 2way ANOVA (genotype/age), of 5-months old WT (n=5), 15-months old WT (n=5), 5-months old Aim2KO (n=6), 15-months old Aim2KO (n=6).). *p<0.05, **p<0.01, ***p<0.001, ****p<0.0001.

### Absence of AIM2 is associated with decrease in cancellous bone in the appendicular and axial skeleton

The assessment of cancellous bone morphology was conducted in two skeletal regions the appendicular (femur) (**Fig. 2**) and the axial (fourth lumbar vertebra, L4) skeleton (**Fig. 3**). AIM2 null females, showed a 35% decrease in the bone volume to total volume (BV/TV) ratio at the femur distal metaphysis at 5 but not 15 months of age (**Fig. 2A**). There was a 55% decrease in bone mineral density (BMD) (**Fig. 2B**) at both 5 and 15 months of age. The reduction in BV/TV in AIM2 null mice was likely due to a decrease in trabecular bone number (Tb.N) (**Fig. 2D**), as trabecular bone thickness (Tb.Th) showed no significant difference between the groups (**Fig. 2C**).

**Figure 2:**
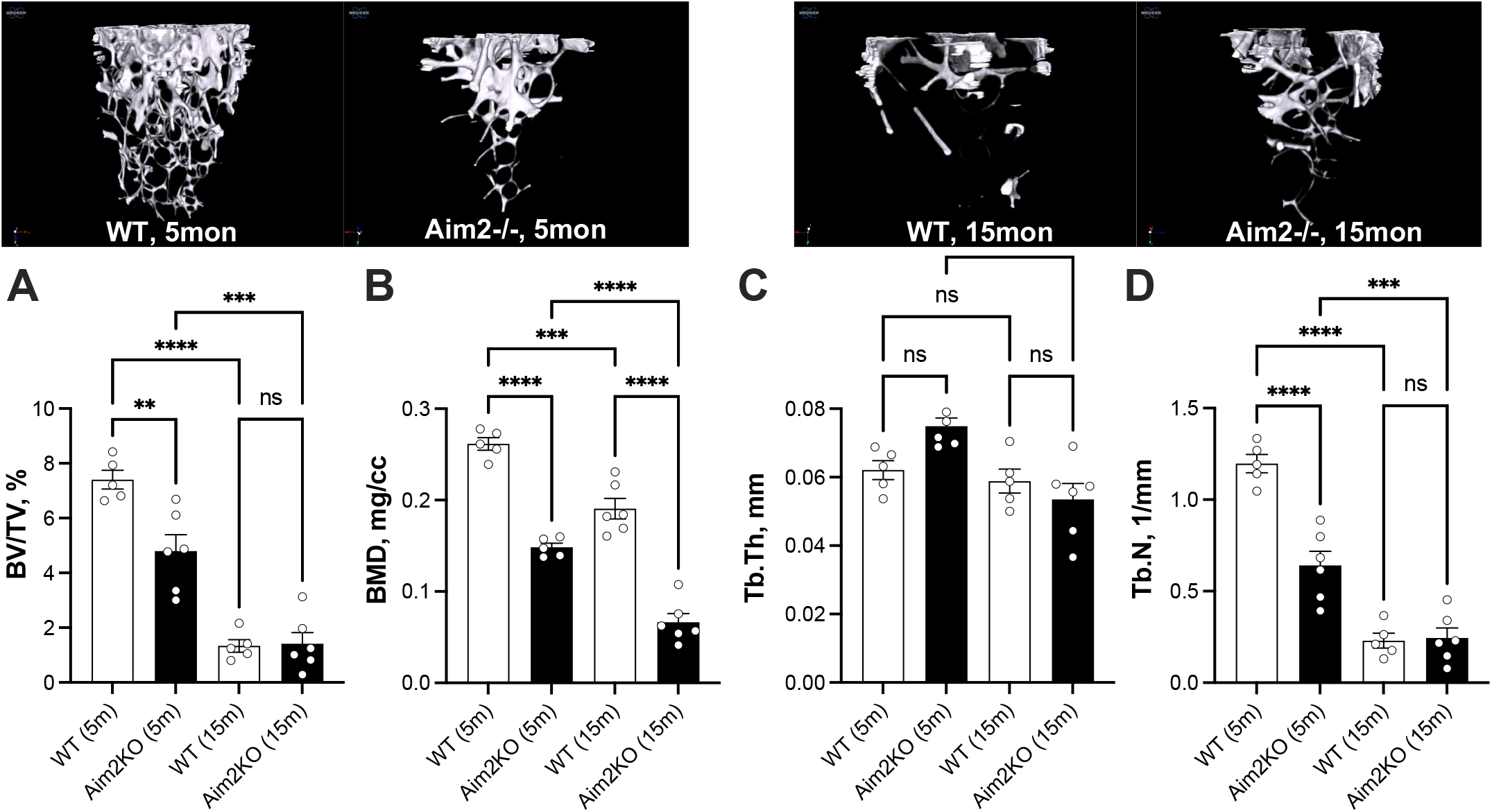
Absence of AIM2 associates with decreased cancellous bone parameters in the appendicular skeleton. Cancellous (trabecular) bone morphology was determined at the femur distal metaphysis including, (A) trabecular bone volume per total volume (BV/TV), (B) bone mineral density (BMD), (C) trabecular bone thickness (Tb.Th), and (D) trabecular number (Tb.N). Significance was determined by 2way ANOVA (genotype/age), of 5-months old WT (n=5), 15-months old WT (n=7), 5-months old Aim2KO (n=6), 15-months old Aim2KO (n=6). **p<0.01, ***p<0.001, ****p<0.0001.

**Figure 3:**
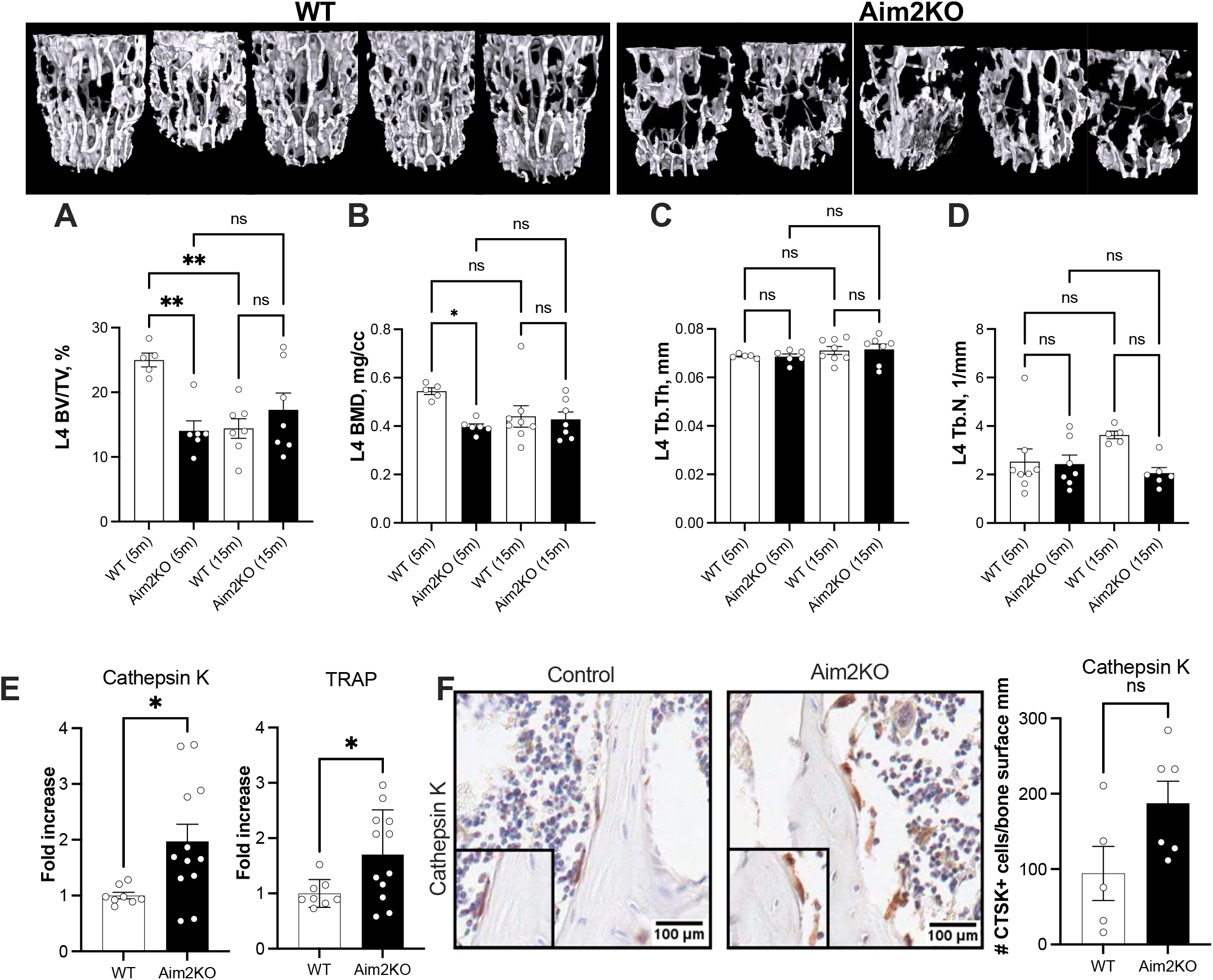
Absence of AIM2 associates with decreased cancellous bone parameters in the axial skeleton. Cancellous (trabecular) bone morphology was determined at the lumbar vertebra-4 (L4) including, (A) trabecular bone volume per total volume (BV/TV), (B) bone mineral density (BMD), (C) trabecular bone thickness (Tb.Th), and (D) trabecular number (Tb.N). 5-months old WT (n=5), 15-months old WT (n=5), 5-months old Aim2KO (n=6), 15-months old Aim2KO (n=7) (E) Cathepsin K and TRAP expression levels were determined in bone marrow of 5-months old mice. WT (n=8), Aim2KO (n=12). (F) Sections of L4 immunostained with cathepsin K antibody. The number of cathepsin K positive cells on bone surface was calculated. WT (n=5), Aim2KO (n=6). Significance was determined by 2way ANOVA (genotype/age).). *p<0.05, **p<0.01.

Similar findings were observed in the trabecular bone morphology of the L4 vertebra, consistent with those observed in the femur distal metaphysis. AIM2 null females displayed a 55% reduction in L4 BV/TV (**Fig. 3A**) and a 30% decrease in BMD (**Fig. 3B**) at 5 months of age, without any changes in Tb.N or Tb.Th (**Fig. 3C, D**). The decreased trabecular number in AIM2 null mice was associated with increased expression of the osteoclast markers Cathepsin K (*Ctsk*) and tartrate resistant acid phosphate (*Trap*) in the bone marrow. Increased levels of Cathepsin K were also detected using immunoreactivity in female AIM2 null mice (**Fig. 3E, F**).

### Absence of AIM2 associates with increased bone marrow adiposity in long bones

In order to gain insights into the bone marrow environment and its potential impact on bone morphology and BMD, we performed H&E staining (**Fig. 4A**), which demonstrated an increase in bone marrow adipose tissue (MAT) in the femurs of AIM2 null mice. We then used flow cytometry analysis of previously described [14] key markers for progenitor of adipocytes and osteochondrogenic cells (**Fig. 4B**). Consistent with the increased adiposity observed throughout the whole body and the bone marrow, we observed an elevation in non-hematopoietic (CD45-), non-endothelial (CD31^−^) cells, expressing the Stem cell antigen (Sca)1 and the surface receptor platelet-derived growth factor-α (CD45-CD31-Sca1+Pdgfa+) cell population in bone marrow indicating increase of progenitor cells with adipogenic potential in 15-month-old female AIM2 null mice. On the other hand, there was a reduction in CD45-CD31-Sca1-Pdgfa+ cell population in the bone marrow reflecting a decrease in the osteochondrogenic cell lineage in both 5 and 15-month-old AIM2 null mice at (**Fig. 4C**). Finally, examination of gene expression in bone marrow samples revealed an upregulation of adipogenic markers such as *Fasn* and *Pparg* in AIM 2 null mice as compared to controls (**Fig. 4D**).

**Figure 4:**
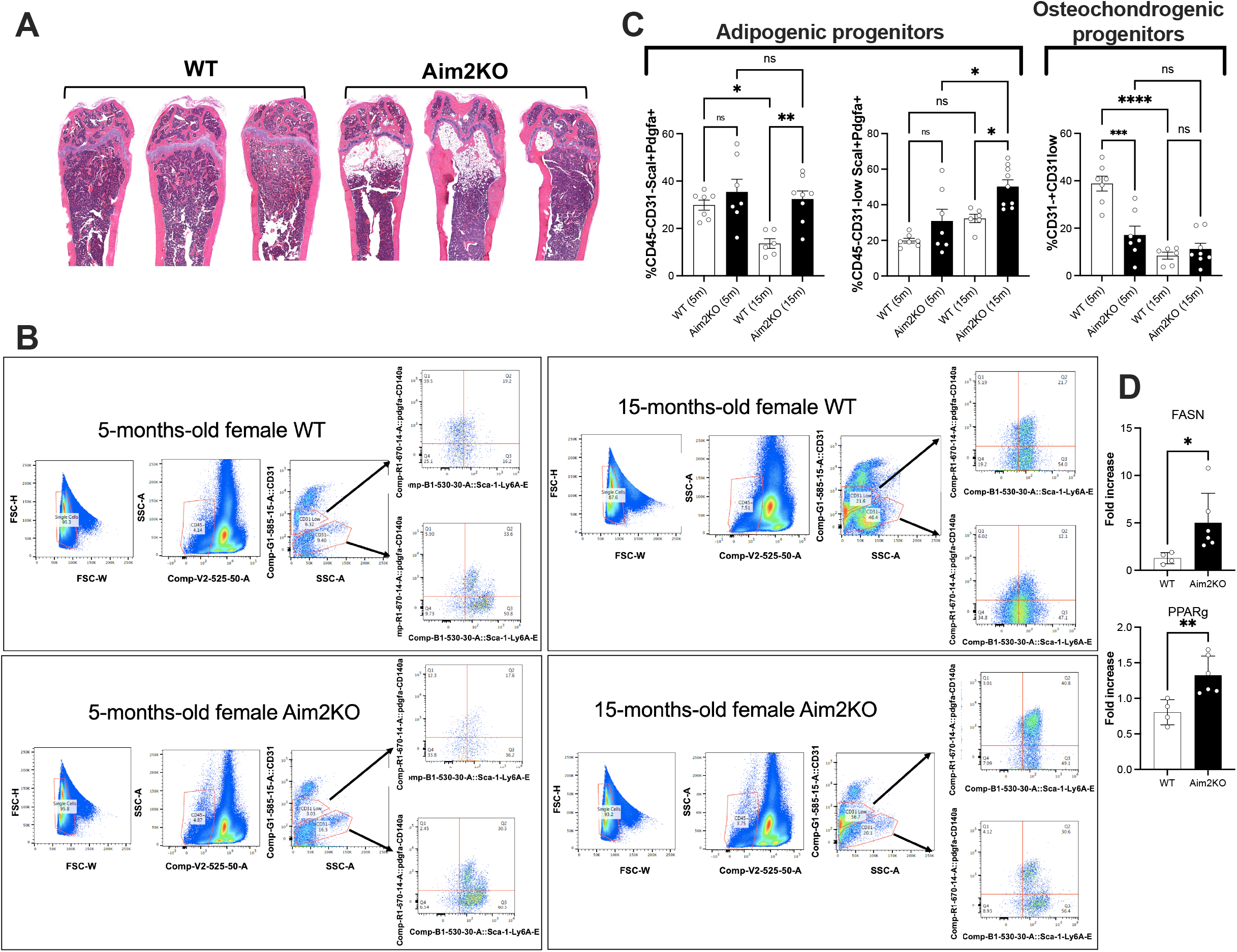
Absence of AIM2 associates with increased bone marrow adiposity in long bones. (A) H&E staining of the distal femur. (B) Representative flow cytometry analyses of bone marrow cells from WT and AIM2 null (AIM2KO) mice at 5 and 15 months of age. Shown are gates for adipogenic progenitor cells CD45-CD31-Sca1+Pdgfa+ and osteochondrogenic CD45-CD31-Sca1-Pdgfa+ cell population in 5 and 15-months-old female WT and AIM2 null mice. First level gating used forward scatter height (FSC-H) versus forward scatter width (FSC-W), followed by side scatter area (SSC-A) versus specific markers (CD31, CD45, Sca1, and Pdgfa). (C) Bar graph demonstrating the flow cytometry analysis. 5-months old WT (n=7), 15-months old WT (n=6), 5-months old Aim2KO (n=7), 15-months old Aim2KO (n=8). (D) Expression levels of the adipogenic markers *Fasn* and *Pparg* in bone marrow was elevated in AIM2 null mice as compared to controls. WT (n=4), Aim2KO (n=6). Significance was determined by 2way ANOVA (genotype/age). *p<0.05, **p<0.01

### Absence of AIM2 is associated with increased inflammation and interferon stimulated genes in the bone marrow

We observed a significant increase in the expression of ifi202b, an interferon-stimulated gene known to regulate adipogenesis in mouse adipose tissue-derived stem cells and promote obesity in mice, in the bone marrow of AIM2 null mice (**Fig. 5A**). The levels of *Ifn-*α*/*β in the bone marrow of 24-month-old AIM2KO mice increased significantly (**Fig. 5B,C**). However, we found no changes in expression levels of *Irf3* gene, a member of the interferon regulatory factor family that plays a crucial role in the induction of type I interferon (IFN-α/β) [22, 23] (**Fig. 5D)**. Notably, however, the total and phosphorylation levels of IRF3 were increased in samples from the AIM2 null mice compared to the controls (**Fig.5E**).

**Figure 5:**
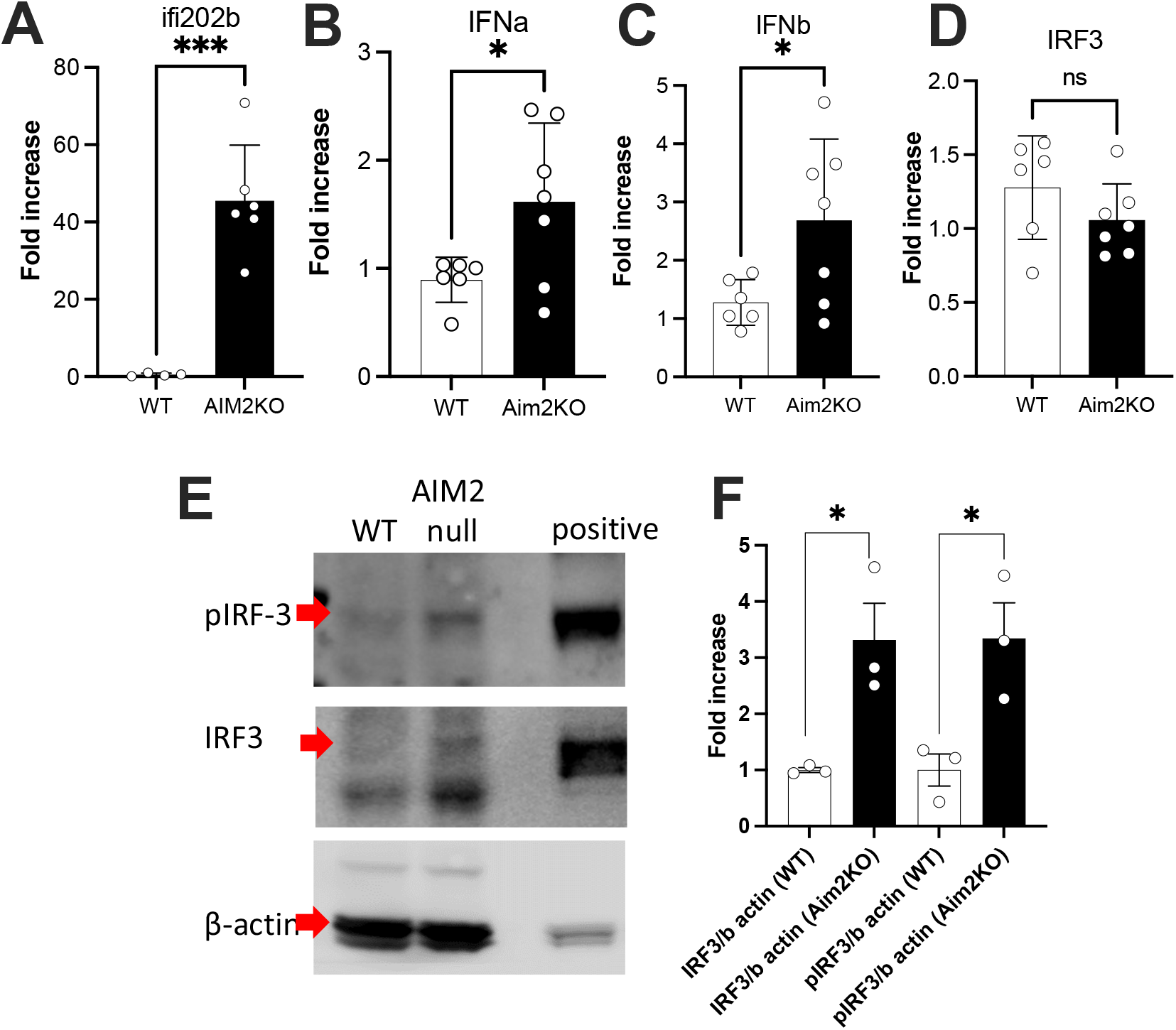
Absence of AIM2 associates with increased expression of inflammation and interferon stimulated genes in bone marrow. Expression of (A) ifi202b, (B) interferon a (IFNa), (C) interferon b (IFNb), and (D) interferon regulatory factor-3 (IRF3) genes in the bone marrow of WT (n=6) and Aim2KO (n=7) mice. (E) Western blot images and (F) quantification of protein extracts from the bone marrow showing the levels of IRF-3 and phosphorylated IRF3 (pIRF3) in WT and AIM2 null mice. WT (n=3), Aim2KO (n=3). Significance was determined by t-test. *p<0.05

## DISCUSSION

In this study, we observed that AIM2 null mice exhibit poor bone mineral density compared to age matched WT control mice. This phenotype is evident at a young age and is associated with increased bone marrow adiposity. We have also identified an increase in markers of adipogenic progenitor cells and a decrease in osteogenic progenitor cells, indicating a shift in the fate of mesenchymal stem cells (MSCs) towards adipogenesis. In addition, we have observed increased expression of osteoclastic markers, which could contribute to compromised bone health.

The increased adiposity in the bone marrow of AIM2 null mice aligns with previous findings of increased marrow adiposity in humans with obesity [15]. Our data suggest a predominance of adipogenic precursors over osteogenic precursors in the bone marrow of AIM2 null mice, indicating a shift in the fate of MSCs towards fibroblasts/preadipocytes at the expense of osteoblasts. Obesity is often associated with systemic inflammation and insulin resistance, which can adversely affect bone mineral density. The adipogenic bone marrow microenvironment also contributes to local inflammation, as indicated by the increase in pro-inflammatory markers in the bone marrow of AIM2 null mice including IFNα and β. It has been shown that DNA damage induces type I IFN production and the increased IFN β promotes cell senescence [16]. Our data indicates that lacking of AIM2 associates with the increased total and phosphorylated IRF3, and expression of IFNα and β in the bone marrow. These pro-inflammatory markers are thought to drive senescence, that potentially can impair bone integrity.

This finding is consistent with previous reports of increased IFN β in immune cells and splenocytes in AIM2 null mice [11]. Increased type 1 interferon signaling leads to the activation of type 1 IFN-stimulated genes (ISGs) and downstream activation of the STAT signaling pathway. Furthermore, our RNA sequencing analysis of white adipose tissue from control and AIM2 null mice revealed that a significant proportion of differentially expressed genes were ISGs, including IFN-inducible p200 family proteins (murine p202 protein encoded by the Ifi202b gene or the human IFI16 gene) [10]. Consistent with increased type 1 IFN signaling and elevated expression of other IFN-inducible genes, such as IRF3 and chemokines CD5L, Cxcl10, and Ccl8, in the adipose tissue of AIM2 null mice [10], we observed increased expression of ifi202b in the bone marrow of AIM2 null mice. Notably, IRF3 has been shown to affect bone health by suppressing osteogenic differentiation of bone marrow mesenchymal stem cells and promoting osteoclast formation, thereby unfavorably affecting bone health [17-19]. Our data showed that although the mRNA levels of IRF3 were not significantly changed in AIM2 null mice at the age of 15-month, the total and phosphorylation levels of IRF3 were increased, indicating a translational and post-translational regulation of IRF3 in AIM2 null mice. Also, the activation of IRF3 in the bone marrow of AIM2 null mice may have an IFN β independent effect on bone health.

The decreased bone mineral density in AIM2 null mice could be attributed to both decreased osteogenesis and increased osteoclastogenesis, as indicated by the presence of increased osteoclastic markers. Additionally, our data suggest a potential link between the cGAS-STING pathway, type 1 IFN signaling, and poor bone health. Increased expression of cGAS and type 1 IFN has been reported in a proportion of patients with systemic lupus erythematosus (SLE), a disease associated with bone loss. Although we have not directly tested activation of the cGAS-STING pathway in the bone marrow, our findings of increased IFN response in AIM2 null mice and its association with poor bone health suggest a potential connection between these activated pathways and bone health in diseases like SLE. The cGAS-STING pathway has been implicated in various inflammatory diseases, including metabolic syndrome, neurodegeneration, myocardial infarction, cardiovascular disease, and fatty liver disease. Therapeutic manipulation of this pathway is being explored to improve immune responses in cancer, viral infections, and chronic inflammatory disorders. Activation of the cGAS pathway has also been linked to senescence. Therefore, the profound changes observed in trabecular bone in our AIM2 null mouse model could be the result of compounding effects from obesity, insulin resistance, inflammation, and exaggerated interferon signaling. Targeting this pathway could offer potential therapeutic approaches to improve bone health in chronic inflammatory conditions such as SLE.

In summary, our findings suggest that AIM2 deficiency associates with poor bone health in mice, likely through promoting adipogenesis. This may be mediated through upregulation of type I IFN signaling and inducing a pro-inflammatory environment, enhanced osteoclast formation that may further contribute to decreased bone mineral density.

## METHODS

### Mice

AIM2 null mice were purchased from The Jackson Laboratory (Bar Harbor, ME, USA) and previously characterized [10]. Aim2 null mice were backcrossed to C57Bl6/J for > seven generations and were housed in the animal facility at the Children’s Hospital of Pittsburgh of UPMC, University of Pittsburgh School of Medicine. Mice (4–5 per cage) were housed under a standard 12 h light/dark cycle (lights on at 07:00) with access to food and water ad libitum. All animal studies were approved by the University of Pittsburgh Institutional Animal Care and Use Committee.

### Micro-computed tomography

Micro-CT was done in accordance with the American Society for Bone and Mineral Research (ASBMR) guidelines [20]. Bones were scanned using a high-resolution SkyScan micro-CT system (SkyScan 1172, Kontich, Belgium) containing 10-M digital detector set at a 10W energy level (100kV and 100 μA), with a 0.5 mm aluminum filter with a 9.7μm image voxel size as described in our recent papers [21-23]. Data reconstruction was done using NRecon software (version 1.7.3.0; Bruker micro-CT, Kontich, Belgium), data analysis was done using CTAn software (version 1.17.7.2+; Bruker micro-CT, Kontich, Belgium) and 3D images were done using CT Vox software (version 3.3.0 r1403; Bruker micro-CT, Kontich, Belgium).

### Histology

Femurs were processed for paraffin embedding and sectioning as described in our recent papers [23]. Sections (7μm) were stained with H&E and TRAP (Sigma, 387A-1KT). Immunohistochemistry was performed as described in our recent papers [23, 24] using antigen unmasking solutionn (H-3300; Vector Laboratories, CA, USA), Cathepsin K (1:200, #ab97110, abcam), HRP conjugated secondary antibody (Cat. No. PK4001, Vector Laboratories, CA, USA), and horseradish peroxidase-3,3′-diaminobenzidine system (cat. no. cts005; R&D Systems, Minneapolis, MN, USA). Images were acquired by Aperio CS2 Scanner (Leica Biosystems, IL, USA) and analyzed by Fiji ImageJ (version 1.51r; NIH, Maryland, USA).

### Flow cytometry

Bone marrow was flushed from femurs, washed and resuspended in FACS buffer (PBS containing 0.5% BSA, 1 mg/ml collagenase IV and 20 μl HEPES). Cell viability was determined in aliquots using trypan blue (Cellgro, Corning, NY, USA). Cells were stained in FACS buffer. CD31 cat#12031182 antibody was purchased from eBioscience (San Diego, CA, USA), CD45 cat#563891, Sca1 cat#557405, and Pdgfa cat#562777 antibodies were purchased from BD Biosciences (San Jose, CA, USA). Adipocytes were identified as CD45-/CD31-/Sca1+/Pdgfa+, osteochondrogenic progenitors were identified as CD45-CD31-Sca1-Pdgfa+. Data acquisition was performed using Fortessa Flow Cytometer (BD Biosciences). Data were analyzed using FlowJo software (Tree Star, Ashland, OR, USA).

### RNA extraction and real-time PCR

BM RNA was extracted using RNeasy purification kit (Qiagen, Valencia, CA, USA) following the manufacturer’s instructions. First strand cDNA was synthesized from 500 ng RNA using iScript cDNA Synthesis Kit (Bio-Rad, Hercules, CA, USA). Real-time PCR was carried out using TaqMan assays in a 10 μl reaction mixture (Bio-Rad) containing 0.1 μl first strand cDNA and 1X probe and primers mix (Bio-Rad). Probes were purchased from Life Technologies or Bio-Rad. Relative mRNA levels were calculated by 2^−ΔΔCt^ and normalized to GAPDH.

### Western Blotting

Bone marrow tissue from WT and AIM2 null mice were homogenized in RIPA buffer (50 mmol/l Tris [pH 7.4], 150 mmol/l NaCl, 1% Triton X-100, 0.5% SDS) containing proteinase inhibitors (Roche, Indianapolis, IN, USA) and phosphatase inhibitors. Total protein (30 μg) was resolved on SDS-PAGE, then transferred to PVDF membranes. The membranes were blocked with 5% non-fat dry milk in Tris-buffered saline (154 mmol/l NaCl) with Tween 20 (TBS-T) for 1 h at room temperature. The membranes were incubated with IRF3 cat#11904 or pIRF3 cat#29047 (abcam, USA) antibodies overnight at 4°C. The membranes were then washed three times with TBS-T and incubated with horse-radish peroxidase (HRP)-conjugated secondary antibodies (diluted in 5% non-fat dry milk in TBS-T) for an additional 1 h at room temperature. Images were taken after adding SuperSignal West Dura extended duration substrate (Thermo Fisher Scientific) to the membrane.

### Statistical analyses

The data are presented as mean ± standard error of the mean (SEM). For the comparison between groups, data were analyzed by two-way analysis of variance (ANOVA) followed by post hoc Tukey’s test (GraphPad Prism version 9.0). The difference between the groups was considered statistically significant when there was p<0.05.

## Acknowledgements

Financial support received from National Institutes of Health Grant R01AG056397 to SY, and S10 OD010751-01A1 for micro-computed tomography.

## Author contribution

ZG followed the mouse cohorts and collected data on body weight, body composition, and performed flow cytometry MD, SBP and GY characterized bone morphology by micro-CT and performed histology. SY and RHM designed and established the mouse cohorts, oversaw all studies involving bone characterizations, summarized data, and prepared the manuscript.

## REFERENCES

1. Camacho, P.M., et al., American Association of Clinical Endocrinologists and American College of Endocrinology Clinical Practice Guidelines for the Diagnosis and Treatment of Postmenopausal Osteoporosis - 2016. Endocr Pract, 2016. 22(Suppl 4): p. 1–42.

2. Hornung, V., et al., AIM2 recognizes cytosolic dsDNA and forms a caspase-1-activating inflammasome with ASC. Nature, 2009. 458(7237): p. 514–8.

3. Choubey, D., et al., Interferon-inducible p200-family proteins as novel sensors of cytoplasmic DNA: role in inflammation and autoimmunity. J Interferon Cytokine Res, 2010. 30(6): p. 371–80.

4. Man, S.M., et al., Critical Role for the DNA Sensor AIM2 in Stem Cell Proliferation and Cancer. Cell, 2015. 162(1): p. 45–58.

5. Wilson, J.E., et al., Inflammasome-independent role of AIM2 in suppressing colon tumorigenesis via DNA-PK and Akt. Nat Med, 2015. 21(8): p. 906–13.

6. Farshchian, M., et al., Tumor cell-specific AIM2 regulates growth and invasion of cutaneous squamous cell carcinoma. Oncotarget, 2017. 8(28): p. 45825–45836.

7. Uresti-Rivera, E.E. and M.H. Garcia-Hernandez, AIM2-inflammasome role in systemic lupus erythematous and rheumatoid arthritis. Autoimmunity, 2022. 55(7): p. 443–454.

8. Colarusso, C., et al., AIM2 Inflammasome Activation Leads to IL-1alpha and TGF-beta Release From Exacerbated Chronic Obstructive Pulmonary Disease-Derived Peripheral Blood Mononuclear Cells. Front Pharmacol, 2019. 10: p. 257.

9. Li, Y.K., J.G. Chen, and F. Wang, The emerging roles of absent in melanoma 2 (AIM2) inflammasome in central nervous system disorders. Neurochem Int, 2021. 149: p. 105122.

10. Gong, Z., et al., Deficiency in AIM2 induces inflammation and adipogenesis in white adipose tissue leading to obesity and insulin resistance. Diabetologia, 2019. 62(12): p. 2325–2339.

11. Panchanathan, R., et al., Aim2 deficiency stimulates the expression of IFN-inducible Ifi202, a lupus susceptibility murine gene within the Nba2 autoimmune susceptibility locus. J Immunol, 2010. 185(12): p. 7385–93.

12. Stadion, M., et al., Increased Ifi202b/IFI16 expression stimulates adipogenesis in mice and humans. Diabetologia, 2018. 61(5): p. 1167–1179.

13. Xin, H., O.M. Pereira-Smith, and D. Choubey, Role of IFI 16 in cellular senescence of human fibroblasts. Oncogene, 2004. 23(37): p. 6209–17.

14. Ambrosi, T.H., et al., Adipocyte Accumulation in the Bone Marrow during Obesity and Aging Impairs Stem Cell-Based Hematopoietic and Bone Regeneration. Cell Stem Cell, 2017. 20(6): p. 771–784 e6.

15. Bredella, M.A., et al., Ectopic and serum lipid levels are positively associated with bone marrow fat in obesity. Radiology, 2013. 269(2): p. 534–41.

16. Yu, Q., et al., DNA-damage-induced type I interferon promotes senescence and inhibits stem cell function. Cell Rep, 2015. 11(5): p. 785–797.

17. Guo, S., et al., Proanthocyanidins attenuate breast cancer-induced bone metastasis by inhibiting Irf-3/c-jun activation. Anticancer Drugs, 2019. 30(10): p. 998–1005.

18. Zhang, Q., et al., Hesperetin Prevents Bone Resorption by Inhibiting RANKL-Induced Osteoclastogenesis and Jnk Mediated Irf-3/c-Jun Activation. Front Pharmacol, 2018. 9: p. 1028.

19. Tsubaki, M., et al., Macrophage inflammatory protein-1alpha induces osteoclast formation by activation of the MEK/ERK/c-Fos pathway and inhibition of the p38MAPK/IRF-3/IFN-beta pathway. J Cell Biochem, 2010. 111(6): p. 1661–72.

20. Bouxsein, M.L., et al., Guidelines for assessment of bone microstructure in rodents using micro-computed tomography. J Bone Miner Res, 2010. 25(7): p. 1468–86.

21. Yildirim, G., et al., Long-term effects of canagliflozin treatment on the skeleton of aged UM-HET3 mice. Geroscience, 2023. 45(3): p. 1933–1951.

22. Poudel, S.B., et al., Excess Growth Hormone Triggers Inflammation-Associated Arthropathy, Subchondral Bone Loss, and Arthralgia. Am J Pathol, 2023. 193(6): p. 829–842.

23. Dixit, M., et al., Induction of somatopause in adult mice compromises bone morphology and exacerbates bone loss during aging. Aging Cell, 2021. 20(12): p. e13505.

24. Poudel, S.B., et al., Sexual dimorphic impact of adult-onset somatopause on life span and age-induced osteoarthritis. Aging Cell, 2021. 20(8): p. e13427.

